# Cyclic parthenogenesis helps populations cross fitness valleys

**DOI:** 10.64898/2026.01.12.699030

**Authors:** Zimo Yang, Nate B Hardy

**Affiliations:** Department of Entomology and Plant Pathology, Auburn University, Auburn, Alabama, USA

**Keywords:** Facultative sex, G-matrix, M-matrix, multivariate fitness landscape

## Abstract

Cyclic parthenogenesis is a form of occasional sexual reproduction. Here, we compare the evolvability of cyclic parthenogens and obligate sexuals in adaptive scenarios that entail the crossing of a fitness valley. We divide the valley crossing process into two phases. In the escape phase, a population produces genotypes that lie on the other side of the fitness valley. In the establishment phase, such genotypes are refined and promoted to high frequency. With individual-based models, we find that although cyclic parthenogens are slower to produce escape genotypes, they more efficiently establish such genotypes, and therefore more rapidly complete fitness-valley crossings. In obligate sexuals, genetic and mutational variance and covariances, **G** and **M,** are readily shaped by selection. In cyclic parthenogens, the shapes of **G** and **M** are less responsive to selection but have relatively little effect on the rate of fitness-valley crossing. In our model, the more adaptive evolution of **G** and **M** in obligate sexuals is not enough to overcome the establishment advantage of cyclic parthenogens. Nevertheless, the relative recalcitrance against selection on **G** and **M** in cyclic parthenogens could be an evolutionary cost that helps explain its paradoxical rarity.

## Introduction

With sign epistasis, an allele that is beneficial when it occurs in one genetic background is deleterious in another. Consequently, the fitness landscape can be “rugged” — it can comprise multiple fitness optima separated by fitness valleys (Weinreich et al. 2005). What determines the evolvability of populations (the ability to respond to directional selection), specifically the ability to overcome sign epistasis and stabilizing selection around a local fitness optimum in a rugged fitness landscape remains an open question (Wright 1932; Wade et al. 2001; Weinreich et al. 2005; Trotter et al. 2014).

The research has proceeded along two major lines. The first has focused on how the storage and release of cryptic genetic variation, a mechanism known as evolutionary capacitance, can expand the neighborhood of evolutionarily-accessible alternative phenotypes and facilitate adaptation (Bergman and Siegal 2003; Trotter et al. 2014; Paaby and Gibson 2016; Zheng et al. 2019). The second line of research has considered how the rate of fitness valley-crossing depends on core population genetic parameters, for example, population structure (Moore and Tonsor 1994) and population size (Weinreich and Chao 2005). Such studies have found that recombination can have strong and complex effects: it promotes the formation of genotypes that lie on the other side of a fitness valley, while also hindering their establishment and fixation (Otto 2009). As a result of this tension, recombination rate can have a non-monotonic effect on the time populations take to cross a fitness valley (Weinreich and Chao 2005; Kim 2007).

In this study, we bridge these research lines by studying how the rate of fitness-valley crossing depends on variation in recombination rate across generations due to alterations between sexual and asexual reproduction. Such a reproduction mode is known as cyclic parthenogenesis, found in many species of agricultural and ecological importance (e.g. aphids, oak gall wasps, walking sticks, and water fleas; Herbert 1987). Theoretical and empirical work suggests that occasional sexual reproduction can be just as good as obligate sexual reproduction for incorporating beneficial mutations, promoting adaptation, and facilitating evolvability, while also incurring fewer of the costs of obligate sexuality (Lynch and Gabriel 1983; Green and Noakes 1995; Yamauchi 1999; D’Souza and Michiels 2010). In particular, cyclical parthenogenesis can reduce recombination load, which may be especially useful in the context of fitness-valley crossing (Weissman et al. 2010). Moreover, cyclic parthenogenesis promotes evolutionary capacitance (Lynch and Gabriel 1983). During parthenogenetic generations, the lack of recombination interferes with selection against mildly deleterious mutations, so new mutations that rescue phenotypes back to the optimum are favored. Groups of compensatory mutations can thus accumulate and increase cryptic genetic diversity. Such cryptic genetic diversity can be released during sexual generations by recombination. As a result, in cyclic parthenogens, the expressed genetic diversity increases sharply in sexual generations and then drops off over subsequent parthenogenetic generations, contrasting with stable equilibrium levels of genetic variance in obligate sexuals (Fig 1 a).

**Figure 1.**
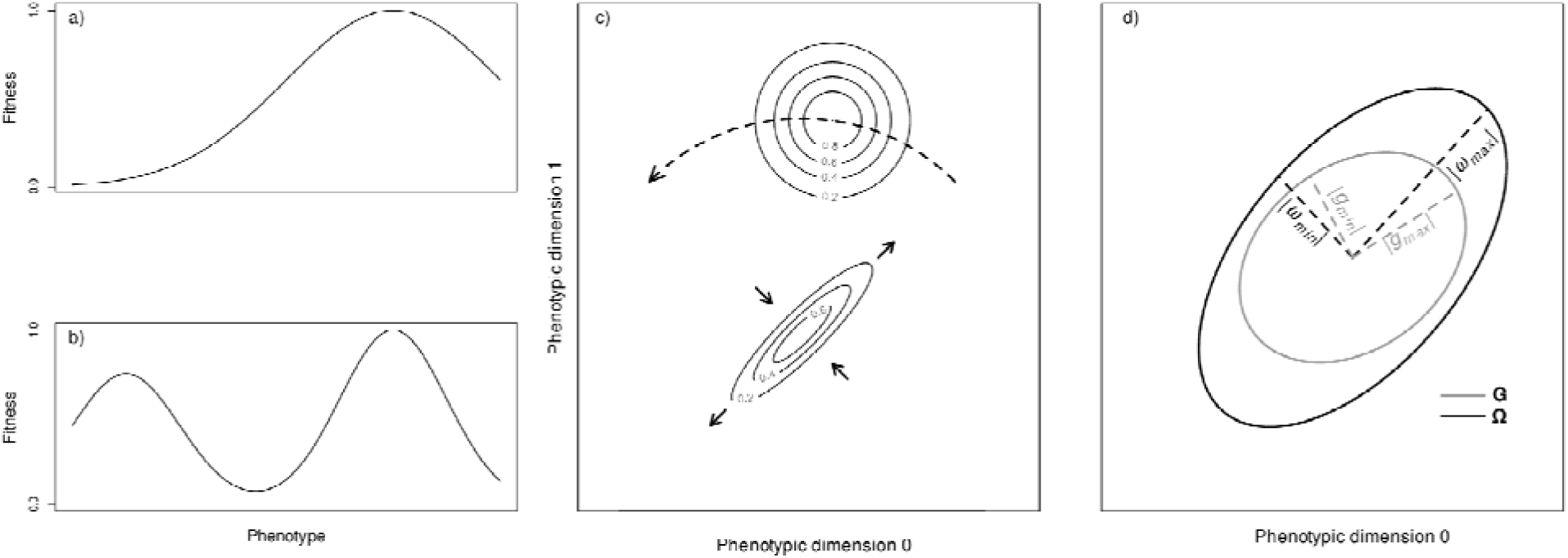
a) A univariate smooth fitness landscape. b) A univariate rugged fitness landscape. In contrast to in a smooth fitness landscape, in a rugged fitness landscape, the fitness effects of mutations that shift the phenotype from the local optimum towards the global optimum are deleterious until the genome accumulates enough mutations to cross the bottom of a fitness valley. **c) A rugged bivariate fitness landscape with a local sub-optimum (lower) and global optimum (upper).** The contour lines are plotted from the modified *Eq. 1* with a strong selection correlation about the local optimum (*r*_ω_ = 0.9) and an intermediate level of alignment between the leading eigenvector of the local fitness optimum and the center of the global fitness optimum (α = 45.0°). Increasing *r*_ω_ of the local optimum stretches it along its leading eigenvector. Increasing α moves the center of the global fitness optimum away from the leading eigenvector of the local optimum. **d) Hypothetical G and** Ω **plotted as 95% confidence ellipses (CE).** For **G**, the ellipse encloses 95% of a population’s phenotypes. For Ω, the ellipse delineates 95% of fitness contour around the local optimum. The dashed lines are the leading (subscript *max*) and second (subscript *min*) eigenvalues plotted in the direction of corresponding eigenvectors. We analyzed **G**-Ω alignment with *r_g_* (the elongation of **G** along |*g_max_*|) and θ (the angle between |*g_max_*| and |ω*_max_*|). **M** is conceptually similar to **G**.

Using individual-based quantitative genetic models, we compare the time to cross fitness valleys between cyclic parthenogens and obligate sexuals across several classes of fitness landscape. First, we consider fitness valleys in univariate phenotypic space without explicit functional epistasis between alleles (Fig. 1 b). To be clear, in this model the phenotypic effect of a mutation is purely additive — there is no functional epistasis (Wade et al. 2001). However, the fitness effect of a mutation depends on its genetic environment — there is statistical sign epistasis. We predict that cyclic parthenogens will cross fitness valleys more readily because 1) the release of accumulated cryptic genetic diversity via periodic recombination allows them to take larger evolutionary steps to escape from the local fitness optimum (evolutionary capacitance) and 2) the parthenogenetic phase of the life cycle will allow escape genotypes to increase in frequency before recombination can break them apart (clonal amplification).

Next, we consider fitness valleys in bivariate fitness landscapes, again with statistical sign epistasis but not functional epistasis. Specifically, we consider a bivariate normal fitness landscape described by a variance-covariance matrix Ω. In this case, adaptive evolution depends on the genetic diversity for each phenotypic dimension as well as the genetic correlation between phenotypic dimensions, which can be described with the genetic variance-covariance matrix **G**. In other words, with more than one phenotypic dimension, we need to account for pleiotropy. With pleiotropy, the genetic correlation between dimensions (*r_g_*) and the leading eigenvector (*g_max_*) of **G** can come to resemble those of the local two-dimensional fitness landscape (*r*_ω_ and *ω_max_*). The value of *g_max_* reflects the direction in the phenotypic space along which most genetic diversity occurs and *r_g_* reflects the relative amount of genetic diversity along this direction (Arnold 2023). When **G** is stable, evolvability is biased towards *g_max_* and increasing *r_g_* magnifies this bias (Schluter 1996; Guillaume and Whitlock 2007). On the other hand, when cryptic diversity is exposed by recombination, it may promote more randomly directed evolutionary steps. Thus, we hypothesize that compared to obligate sexuals, the valley-crossing rate of cyclic parthenogens will be less sensitive to the specific configuration of valleys and optima in the fitness landscape (Fig. 1 c).

Lastly, we consider a bivariate phenotype model with, functional epistasis (Wade et al. 2001). Although the cause of ruggedness in the fitness landscape (i.e. the phenotype-fitness map) is still sign epistasis, we let the contribution of each allele to the phenotype dependent on its genetic environment. With functional epistasis and pleiotropy, we expect an expanded scope for selection to shape **G**, as well as **M**, the variance-covariance matrix of mutational effects. Selection can shape the loading of additive genetic effects across epistatic loci so that ― even without changing the mutation process itself ― the mean mutation load across loci can be reduced. Thus, **M** as well as **G** can align with Ω (Jones et al. 2014). It is unclear how cyclic parthenogenesis affects this alignment, but any such effects could indirectly influence evolvability. Specifically, **M**-Ω alignment can stabilize **G** around Ω and strengthen the effect of **G** on evolutionary trajectory (Jones et al. 2007). This effect might be weaker in cyclic parthenogens than in obligate sexuals, as each sexual generation releases cryptic genetic diversity that can scramble the loading of additive genetic effects across loci. Thus, we hypothesize that with epistasis, cyclic parthenogens will not evolve **M** to align with Ω as efficiently as obligate sexuals. If this holds, on fitness landscapes with eccentric local optima (high *r*_ω_) and strong alignment between *ω_max_* and the global fitness optimum, any evolvability advantage for cyclic parthenogenesis could be diminished.

## Methods

### Univariate phenotype without epistasis

We modeled the evolution of cyclic parthenogenetic and hermaphroditic populations that undergo one sexual generation every *ρ* generations. This setup most closely resembles cyclic parthenogenesis in digenean (Digenea: Trematoda; Herbert 1987). Analyzing hermaphroditic instead of gonochoric populations allows us to focus on the evolutionary consequence of recombination per se without other, potentially confounding effects of sexual reproduction (Otto 2009).

Each population is panmictic and has a carrying capacity, *K*. Individuals are diploid with one univariate quantitative phenotype determined by *L* loci. The first event in each generation is reproduction, either via cloning or random mating. Each parent has a litter size sampled from a Poisson distribution with an expectation of two. Mutation occurs at a rate *µ* = 0.008 per locus per gamete. When a mutation happens at a particular locus, an effect sampled from a normal distribution with a mean of zero and standard deviation of *m* is added to the existing genetic value of that locus. The mutational process thus creates an influx of total genetic variance *V_m_ = 2Lµm^2^* per generation per offspring. The genotype of an individual is the sum of all genetic values across loci, and the phenotype is the genotype value pulse environmental noise sampled from a normal distribution with a mean of zero and a variance of *V_n_*. During sexual reproduction, recombination takes place with a breakpoint occurring between two adjacent loci at rate *R*.

Selection happens after reproduction. An individual’s fitness is then calculated based on the matching between its phenotype and phenotypic optima:

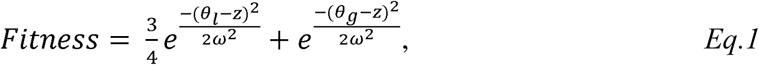

where *θ_l_* and *θ_g_* are respectively the local and global optimal phenotypes, *z* is the phenotype of the individual, *ω* is the weakness of selection. Fitness is then scaled by *K*/*N* where *N* is the population size; in other words, a component of fitness is determined by negative density dependence. An individual’s fitness determines their probability of surviving to the next generation. Overlapping generations are permitted.

### Bivariate phenotype without epistasis

This model extends the univariate phenotype model by letting all mutations pleiotropically affect two phenotypic dimensions. Each mutation’s genetic effects are sampled from a bivariate normal distribution with means of zero, standard deviations of *m*, and a correlation coefficient of zero. As in the univariate model, a mutation’s effect on each phenotype dimension is added to the current genetic values at a locus. The genotype value and phenotype of each dimension are calculated in the same way as for the univariate phenotype. Fitness is calculated by substituting a bivariate normal PDF into *Eq. 1*. Specifically, *z* and *θ_i_* in *Eq. 1* are replaced with row matrices **z** and θ_i_ to account for the two phenotypic values, and each *ω_i_* term in Eq. 1 is replaced with a corresponding selection variance-covariance matrix:

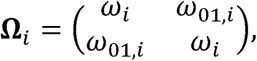

where *ω_i_* is the weakness of direct selection on each phenotypic dimensions (we used the same strength on both dimensions) and *ω_01,i_* is selection covariance. Then *r*_ω*,i*_, the selection correlation, is *ω_01,i_ / ω_i_*. The same kind of selection matrix is used to describe both the local and global fitness optima, but in our analysis, the Ω of the global optimum is fixed whereas that of the local optimum varies. For convenience, from here on we describe the fitness landscape by using the parameters Ω, *r*_ω_, and *ω_max_* without qualifying that these refer to the local optimum.

To explore the effect of the shape of the fitness landscape (Ω) on **G** — and consequently the effect of Ω on the rate of fitness-valley crossing — we ran simulations with 1) the *r*_ω_ of the local optimum varying from 0.0 to 0.9 (Fig. 1 c), and 2) *α*, the angle between the center of the global fitness optimum and *ω_max_*varying from 0° to 90° (Fig. 1 c). During these manipulations, the strength of direct selection on each phenotype and the Euclidean distance between the centers of local and global fitness optima were kept constant.

### Bivariate phenotype with functional epistasis

Further extending the bivariate phenotype model to include functional epistasis brings us to the multi-linear genotype-phenotype map used in Jones et al. (2014). Phenotypes are now determined as follows:

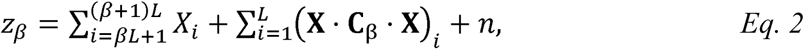

where *z*_β_ is the value of phenotypic dimension *β* (note that the two dimensionalities are respectively denoted as 0 and 1), **X** is a 1-by-2*L* row matrix of genotypic values, *X_i_* is the *i*th element in the row of **X**, **C**_β_ is a matrix of epistasis coefficients for phenotypic dimension *β*, and *n* is environmental noise. *X_1_*to *X_L_* store the genetic values for phenotypic dimension 0; the sum of these values is the additive genetic effect of all alleles on phenotypic dimension 0. Similarly, *X_(L+1)_* to *X_2L_*are used for phenotypic dimension 1. We allowed for epistasis between all pairwise combinations of loci. The epistatic coefficients are stored in **C**_β_, a 2*L*-by-2*L* matrix. The elements of the upper triangle of **C**_β_ are zeros so that each epistatic interaction effect is only represented (and counted) once. The diagonal elements of **C**_β_ and the diagonal elements of the sub-matrix **C**_β_[{*L*+1, 2*L*}, {1, *L*}] are also set to zero so that there is no interaction between a locus and itself. Therefore, the dot product **X** *·* **C**_β_ *·* **X**^T^ is a 1-by-1 matrix whose only element is the total epistatic effect of all interacting genetic values on phenotypic dimension *β*. In other words, the epistatic effect on a phenotypic dimension is the sum of the products of all additive genetic value pairs and their interaction coefficients. Following Jones et al. (2014), the network of pairwise epistatic interaction coefficients is a population-level model parameter, with each coefficient sampled once at the start of each simulation from a normal distribution with a mean of zero and standard deviation of *V_e_*.

### Simulation

Simulations were run with a modified version of SLiM 5.1 (Haller et al. 2026); we added a function to efficiently calculate total genetic diversity (https://zenodo.org/records/20642543). Results were analyzed with R 4.5.1 (R Core Team 2025). Model parameters are summarized in Table 1 and model codes are provided in as a Zenodo repository (https://zenodo.org/records/20642543). Each simulation starts with a monomorphic population perfectly adapted to the local phenotypic optimum (the first term of *Eq. 1* only) passing through 10,000 generations to ensure that mutation-selection-drift balance is attained. We ran two sets of experiments. First, to validate the model, we tried to recover the major findings of Lynch and Gabriel (1983) that although cyclic parthenogenesis induces pulses of expressed genetic diversity, it does not affect the overall rate at which populations respond to directional selection on a smooth fitness landscape. For this experiment, after the first 10,000 generations, the local fitness optimum was displaced by five phenotypic units for a univariate phenotype or four units for a bivariate phenotype (slightly less than for the univariate model to avoid extinction). We then monitor changes in mean population fitness over the following 300 generations.

**Table 1.** Key model parameters and values explored.

| Parameter | Symbol | Core Values | Range explored |
| --- | --- | --- | --- |
| Carrying capacity | $K$ | 1000 | 1000 |
| Number of loci | $L$ | 20 | 8-20 |
| Frequency of sexual reproduction | $\rho$ | [1, 10] | [1, 10] |
| Recombination rate | $R$ | 0.001 | 0.0001-0.0100 |
| Variance of additive mutational effects | $V_m$ | 1 | 0.0001-1.0000 |
| SD of epistatic coefficients | $V_e$ | 0.2 | 0.2 |
| SD of environmental noise | $V_n$ | 0.5 | 0.1-2.0 |
| Weakness of direct selection about the local optimum* | $\omega$ | 2 | 2.0-49.0 |
| Selection correlation in the about the local optimum | $r_\omega$ | 0 | 0.0-0.9 |
| Angle between optima | $\alpha$ | 0 | 0°-90° |

\*Same value for both phenotypes for both optima.

In the second set of experiments, to explore the effect of cyclic parthenogenesis on the rate of fitness-valley crossing, at the 10,001^th^ generation we introduce a second, higher optimum six phenotypic units (five for bivariate phenotypes) away from the local optimum. (All threshold values were experimentally determined so that populations can consistently but not instantly cross the fitness valley.) Simulations then continue until 40,000 generations have elapsed, or until the population crosses the fitness valley with a mean phenotype close to the global optimum. For the univariate phenotype model, our threshold is one phenotypic unit from the global optimum, and two for the bivariate phenotype models. We marked this time to cross the valley as T_tol_. We also tracked T_esc_, the time populations take to generate at least one individual that escapes the local optimum. We define an escape phenotype as occurring within half of the Euclidean distance from the local to the global fitness optimum. Finally, the time populations taken to establish around the global optimum, T_est_, is T_tol_ – T_esc_.

To better understand the dynamics of each experiment, we characterized the effect of cyclic parthenogenesis on **G** and **M** before the manipulation of fitness landscape in experiment 2, specifically, at every generation from the 9,900^th^ to 10,000^th^. Details of this characterization are provided in the next section.

### Assays for **G**, **M**, and cryptic genetic diversity

Based on the way in which cyclic parthenogens accumulate and release cryptic genetic diversity, we predicted that in comparison to obligate sexuals (1) their phenotypic evolution would be less restricted by **G** and (2) there would be less alignment between **M** and Ω.

We estimated **G** via the mid-parental regression as described in (Falconer and Macky 1996; Phillips et al. 2001). In brief, each individual was crossed with another randomly chosen individual and produced two offspring. The diagonal elements of **G** were estimated as twice the covariance between mid-paternal and offspring for the corresponding phenotypic dimension. The off-diagonal elements of **G** were estimated as the sum of the covariance between mid-parental phenotypic dimension 0 and offspring phenotypic dimension 1 and the covariance between mid-paternal phenotypic dimension 1 and offspring phenotypic dimension 0.

We estimated **M** with a mutate-and-re-phenotype assay, following the approach of Jones et al. (2014). This assay was run on 100 randomly sampled individuals. For each locus, 50 mutations were generated with additive allele effects sampled from the same bivariate normal distribution that was used in the main flow of the simulation. For each of these mutations, for each genome of each sampled individual, we measured the change in each phenotypic dimension. To be clear, this accounts for a mutation’s additive and epistasic effects. But by averaging the phenotypic deviation caused by a mutation across genetic backgrounds sampled according to their frequency, we arrive at a characterization of additive mutational variance. So, a locus-specific **M** was estimated for all 50 mutations and 100 individuals. We then estimated the genome-wide **M** as the mean of the locus-specific estimate of **M**.

To characterize the evolutionary changes in **G** and **M**, we calculated two metrics, which were calculated in the same way for **G** and **M** so we only describe the calculations for **G**. 1) To measure to what degree the genotypic diversity are allocated around the direction of less fitness cost, we measured the orientation of **G** as the angle between **G**’s leading eigenvector (*g_max_*) and that of Ω. 2) To track how much genetic diversity distribution around *g_max_*, We measured the shape of **G** as the genetic correlation, *r_g_*. 2These metrics are illustrated in Figure 1 d.

Cyclic parthenogens accumulate cryptic genetic diversity as compensatory pairs of genetic values. Lynch and Gabriel (1983) analytically calculated cryptic genetic variance from the covariance among alleles, but this approach would be computationally prohibitive for our models. Instead, we indirectly tracked cryptic genetic diversity by looking at parent-offspring resemblance, using the same mid-parental regression as for the diagonal elements of **G**. To be clear, resemblance varies according to the amount of genotypic diversity (Falconer and Macky 1996). We measured the resemblance of both the genotype value (the sum of genetic values; denotated *h^2^*) and the absolute genotype value (the sum of the absolute values of genetic values, which accounts for the amount of compensatory genetic values; denotated |*h^2^|*). During fitness valley-crossing, with repeated production of escape genotypes via the exposing of cryptic genetic diversity, *h^2^* should temporally increase and |*h^2^|* should temporally decrease.

## Results and Discussion

### Model validation

Under directional selection along a smooth fitness landscape, our individual-based model behaves similarly to a modified Lynch-Gabriel analytical model (https://zenodo.org/records/20642543) that scales genetic variance in each generation by a fixed factor of linkage disequilibrium (Bulmer 1974; Lande 1975). The population mean phenotype evolves towards the fitness optimum at a similar rate in obligate sexuals and cyclic parthenogens, although in cyclic parthenogens the phenotypic variance rises and falls in synchrony with the alteration of reproduction modes (Fig. 2 a & b). One discrepancy between our individual-based model and the analytical model is that at the beginning of directional selection the individual-based model generates higher phenotypic variance (Fig. 2 a). This is likely due to initial deviations of genetic diversity from a normal distribution, which violates the assumption of the analytical model (Lynch and Gabriel 1983).

**Figure 2.**
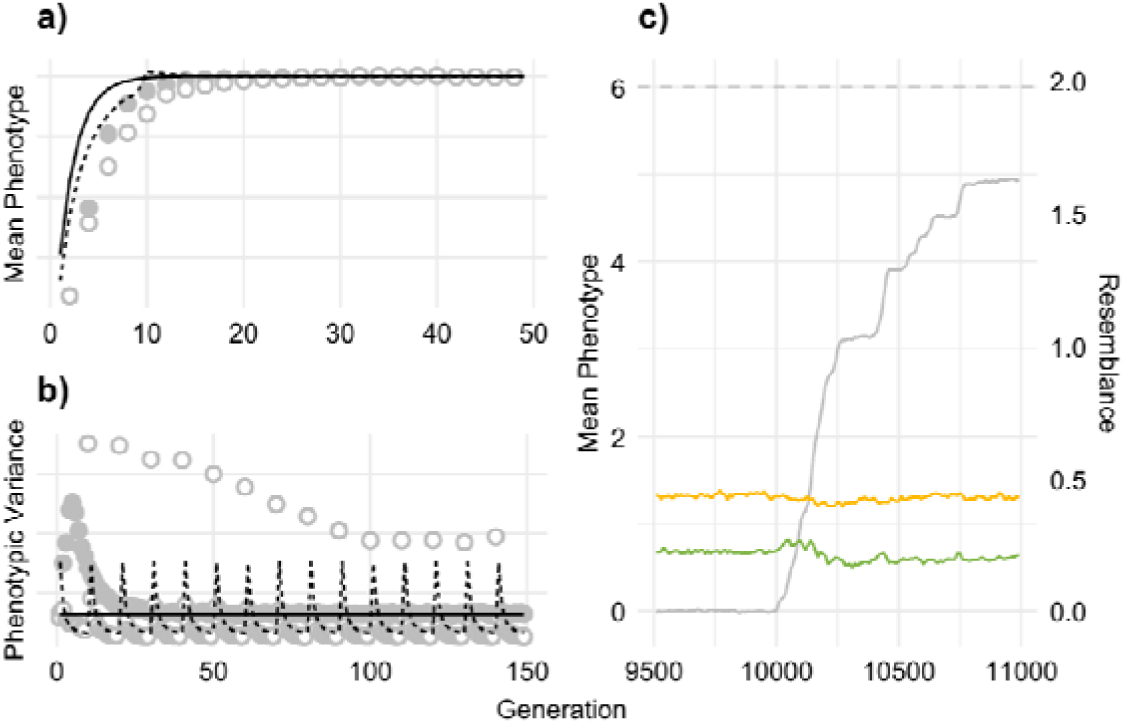
Model validation. Comparison of dynamics of a) the mean phenotype and b) phenotypic variance under directional selection on a smooth fitness landscape under the simulation and analytical models. Lines: analytical solution of the Lynch-Gabriel model; points: mean values of 30 replicates of the simulation. Solid lines and points: obligate sexuals; dashed lines and hollow points: cyclic parthenogens. Directional selection starts at generation 0. The mean phenotype reaches the equilibrium at a similar rate between models and reproduction mode. The simulated phenotypic variance is higher than the analytical variance at the start of directional selection but quickly aligns. **c) Parent-offspring genetic resemblance changes at the beginning of fitness valley-crossing in cyclic parthenogens.** Data are measurements every 50 generations averaged across 30 replicates with the univariate phenotype and core parameter set. Dashed line: the global fitness optimum. Green: the resemblance between parents and offspring of the sum of genetic values (*h^2^*); yellow: the resemblance between parents and offspring of the sum of absolute values of genetic values (|*h^2^*|); grey: mean phenotype. The global fitness optimum is added at generation 10, 000. Immediately after the addition of the global fitness optimum and around the onset of mean phenotype increases, *h^2^* increases while |*h^2^*| slightly drops.

The individual-based model supports that the release of cryptic genetic diversity contributes to the escape stage of fitness valley-crossing in cyclic parthenogens. When a population’s mean phenotype begins to move towards the global optimum, the resemblance of genotypic values (the sum of genetic values) between parents and offspring (*h^2^*) increases while the resemblance of the sum of absolute genetic values between parents and offspring (|*h^2^*|) drops (Fig. 2 c). This is evolutionary capacitance in action; cryptic genetic variance is converted to additive genetic variance that fuels fitness valley crossing. Obligate sexuals — in which some cryptic genetic diversity can accumulate via compensatory mutations between tightly-linked loci — display similar but weaker dynamics (Fig. S1).

### Cyclic parthenogenesis helps populations cross univariate fitness valleys

To reiterate, following Weissman et al. (2010), we separated fitness valley-crossing into two processes. The population first needs to generate at least one individual with a phenotype that escapes the basin of attraction around the local fitness optimum. The time to achieve this is denoted as T_esc_. Then, the escaped phenotype needs to be refined by selection and increase in frequency. We call this process ‘establishment’ and denoted the time it takes as T_est_. The total time it takes to cross a fitness valley is then T_tol_ = T_esc_ + T_est_.

Cyclic parthenogens tend to have larger T_esc_ than obligate sexuals, unless escape is instantaneous regardless of reproduction mode (Fig. 3 a). Even though cyclic parthenogens produce pulses of expressed genetic diversity and have the potential to take larger evolutionary steps at each sexual generation (the evolutionary capacitance effect; Lynch and Gabriel 1983; Fig. 2 b), such steps are too seldom taken to escape the local optimum as efficiently as with the smaller-on-average yet more frequent steps taken by obligate sexuals. On the other hand, compared to obligate sexuals, cyclic parthenogens have smaller T_est_ (Fig. 3 b). This is consistent with clonal amplification causing a reduction in recombination load. This inference is further supported by reproduction-mode-dependent responses to variation in recombination rate (*R*). In obligate sexuals, T_est_ is lowest at intermediate *R,* which balances the odds of forming and breaking escape genotype (Fig 3 b; Weissman et al. 2010). In contrast, in cyclic parthenogens, T_est_ monotonically declines as *R* increases (Fig. 3 b) — parthenogenesis shelters newly-formed escape genotypes from recombination, while increasing *R* increases the efficiency of the conversion of cryptic genetic variance into additive genetic variance. The T_tol_ pattern is identical to that of T_est_ as T_est_ >> T_esc_ (cyclic parthenogens: T_esc_ ∈ [1, 20], T_est_ ≥ 33; obligate sexuals: T_esc_ [1, 17], T_est_ ≥ 108).

**Figure 3.**
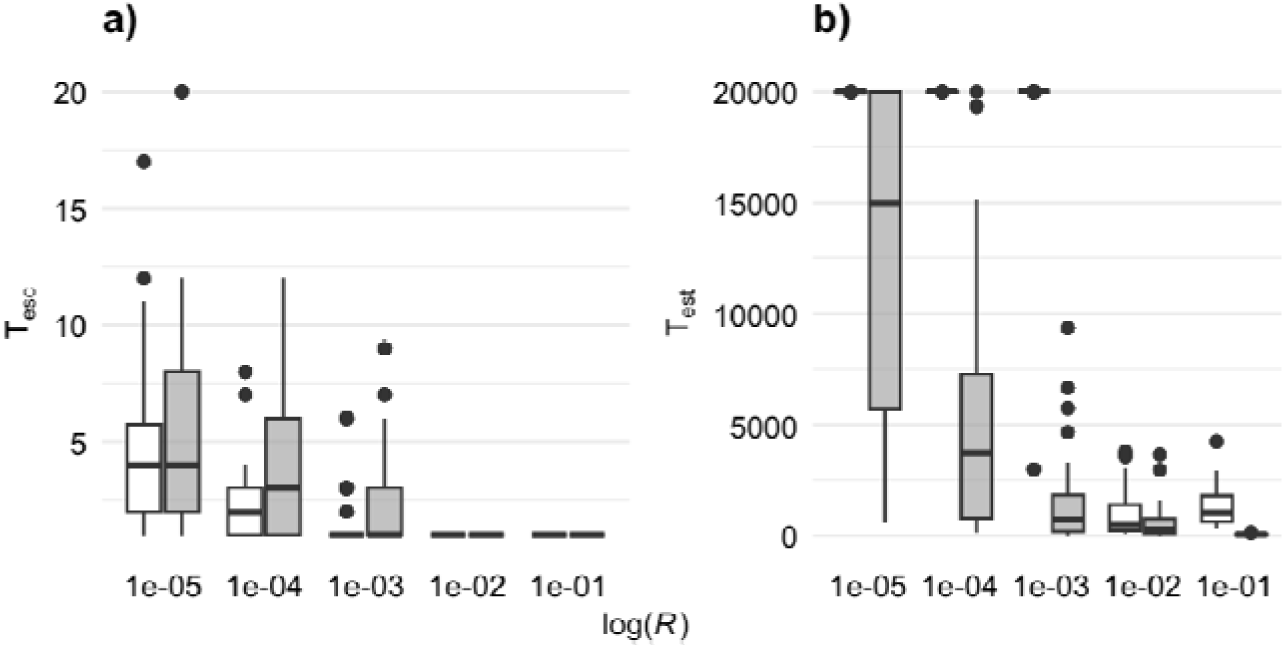
For the univariate phenotype, how recombination rate affects a) the time taken to escape from the local fitness optimum and b) the time taken to establish around the global fitness optimum. White: obligate sexuals; grey: cyclic parthenogens. The bottom, center, and top line of each box mark the 1^st^, 2^nd^, and 3^rd^ quantiles respectively. Whiskers show 1.5 times the inter quantile distance. Each bar is from 30 replicates with core parameters. Cyclic parthenogens have larger T_esc_ but lower T_est_ than obligate sexuals. Cyclic parthenogenetic T_est_ monotonically decreases as R increases; this contrasts with the convex mapping of *R* to T_est_ in obligate sexuals.

The tendency for cyclic parthenogens to have larger T_esc_ but lower T_est_ than obligate sexuals holds across variation in mutational influx per generation (*V_m_*), genome size (*L*), and environmental noise (*V_n_*) (Fig. S2). The only exception is that in cyclic parthenogens T_est_ becomes similar or slightly larger than in obligate sexuals when *V_n_* is large enough to completely overshadow the fitness valley effect. In general, T_esc_ and T_est_ are diminished by increasing *V_m_*, *L*, and *V_n_* (Fig. S2) — increasing *V_m_* and *L* increase additive genetic variance, while larger *V_n_* reduces the efficiency of stabilizing selection around the local fitness optimum.

### Phenotypic valley-crossing of cyclic parthenogens is less sensitive to **G**

The genetic variances and covariance of multivariate phenotypes can be shaped by selection (Steppan et al. 2002). Our bivariate, additive phenotype model shows that in comparison to obligate sexuals, cyclic parthenogens evolve **G** with leading eigenvectors less aligned with Ω (Fig. 4 a), although the matching of *r_g_* to *r*_ω_ is consistent across reproductive modes (Fig. 4 b). Visualizations of **G**s in 95% confidence ellipse are in Fig. S3. In other words, compared to obligate sexuals, cyclic parthenogens evolve genetic correlations of similar strength, but the direction of such correlations in the phenotypic space is less stable, and hence aligns more poorly on average to the shape of the selective landscape. This is consistent with the hypothesis that in cyclical parthenogens the efficiency of selection on **G** is reduced by pulses of released cryptic genetic variance.

**Figure 4.**
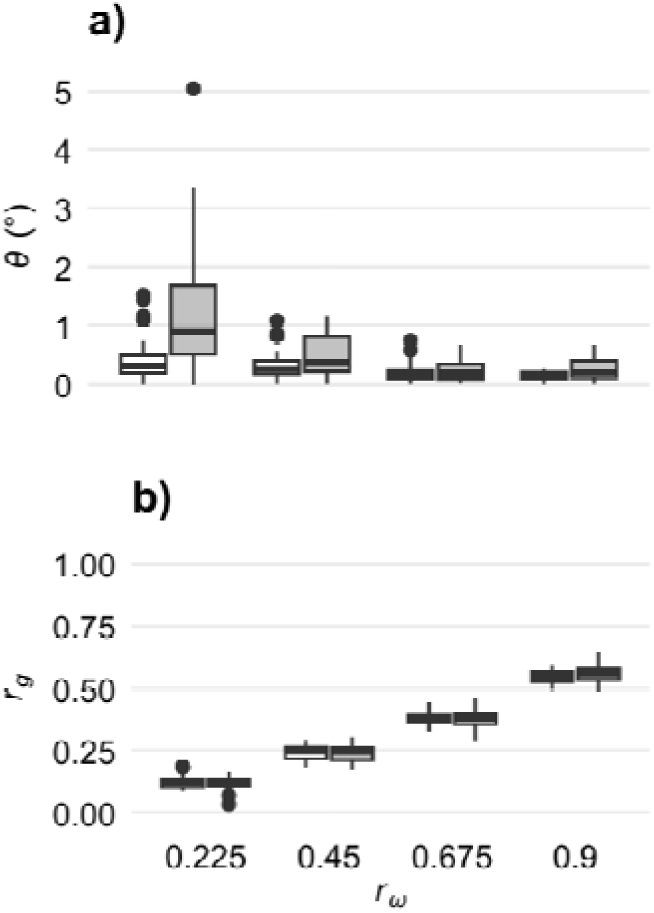
G-Ω alignment in terms of a) the direction of leading eigenvectors and b) the genetic correlation for a bivariate, additive phenotype. White: obligate sexuals; grey: cyclic parthenogens. The bottom, center, and top line of each box mark the 1^st^, 2^nd^, and 3^rd^ quantiles respectively. Whiskers show 1.5 times the inter quantile distance. Each bar is from 30 replicates with the core parameter set. Cyclic parthenogens tend to have poorer alignment of G with Ω (larger θ) than obligate sexuals but the difference diminishes as *r*_ω_ increases. Cyclic parthenogens and obligate sexuals have similar *r_g_* across values of *r*_ω_.

In line with previous work showing that phenotypic evolution is biased towards the direction of **G**’s leading eigenvector, *g_max_*, we found that in obligate sexuals T_est_ tends to be smaller when *r*_ω_ is large (and consequently so is *r_g_*) and the global optimum lies closer to *g_max_* (_α_ is small; Fig. 5; Schluter 1996; Guillaume and Whitlock 2007). On the other hand, in cyclical parthenogens, we found that _α_ has little effect on the rate of fitness-valley crossing (Fig. 5 b). This is consistent with our prediction that in cyclic parthenogens fitness valley-crossing will depend more on non-directional evolutionary capacitance and clonal amplification, and therefore be less sensitive to the shape of **G**. Even when the local optimum is strongly eccentric and well-aligned with the global optimum, cyclic parthenogens still have smaller T_est_ (Fig. 5). When a fitness-valley is a formidable barrier, T_est_ ≈ T_tol_, and T_esc_ — which is more sensitive to the local contours of the selective landscape — has little effect on the overall rate of valley crossing (Fig. S4).

**Figure 5.**
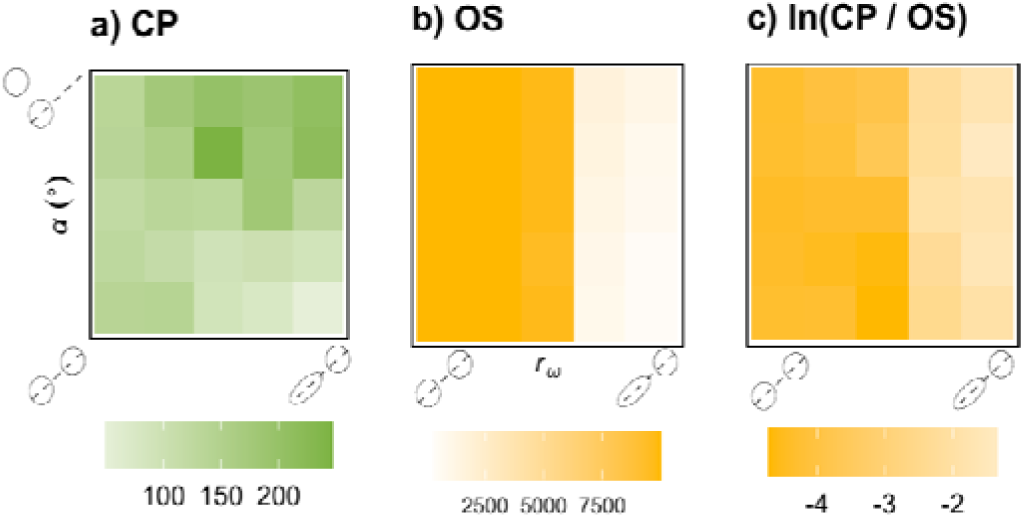
T_est_ across fitness landscapes of various shapes in a) cyclic parthenogens, and b) obligate sexuals, along with c) the log-ratio of T_est_ in cyclic parthenogens over obligate sexuals. Each block is the average of 30 replicates. Five evenly distanced values across [0.0, 0.9] were used for *r*_ω_ and across [0, 90°] for α. For cyclic parthenogens, small α and small *r*_ω_ minimize T_est_ while large α and small *r*_ω_ maximize it. For obligate sexuals, *r*_ω_ has a strong negative effect on T_est_, overwhelming the effect of α. Obligate sexual T_est_ is always larger than cyclic parthenogenetic T_est_.

### Cyclic parthenogenesis has little effect on M

Previous work has shown that with functional epistasis, selection can shape the mutational variance and covariance, **M**, so as to diminish mutation load (Jones et al. 2014). We found that this effect is more pronounced in obligate sexuals than in cyclic parthenogens (Fig. 6 b & S5). To reiterate, in the multilinear model of functional epistasis, each simulation takes place under a matrix of pairwise epistatic interaction coefficients that are sampled from a normal distribution at the start and then fixed. **M** evolves via changes in the loading of genetic values across loci, which is a difficult and slow optimization process (Jones et al. 2014). Weaker cyclic parthenogenetic **M**-Ω alignment is an indication that **M** evolution is easier with relatively low and steady levels of additive genetic variance than with periodical pulses of high genetic variance. Previous work has also shown that **M** can stabilize **G** (Jones et al. 2007), and indeed, we find that cyclic parthenogens also have poorer **G**-Ω alignment — weaker alignment of the leading eigenvectors and less stable genetic correlation compared to obligate sexuals (Fig. 6 a). However, this could also be due in part to functional epistasis per ser, which provides another channel for the accumulation and release of cryptic genetic variance. As for the bivariate phenotype model without functional epistasis, the rate of fitness valley-crossing in cyclic parthenogens is less sensitive to **G-**Ω alignment but still faster than in obligate sexuals, regardless of the shape of the selective landscape (Fig. S6).

**Figure 6.**
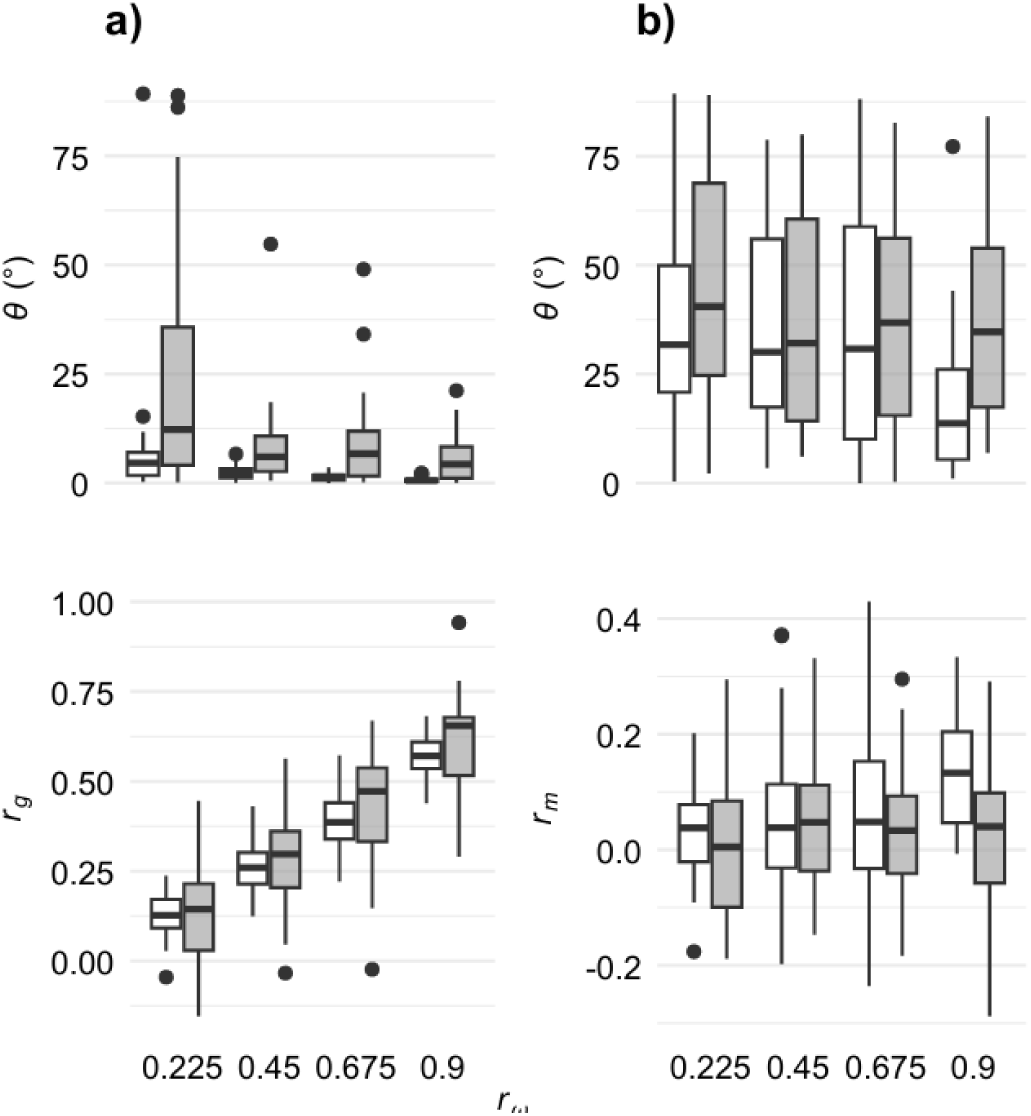
a) G-Ω alignment and b) M-Ω alignment in the bivariate phenotype model with functional epistasis. White: obligate sexuals; grey: cyclic parthenogens. The bottom, center, and top line of each box mark the 1^st^, 2^nd^, and 3^rd^ quantiles respectively. Whiskers show 1.5 times the inter quantile distance. Each bar is from 30 replicates with the core parameter set. When *r*_ω_ is large, in cyclic parthenogens, **M** tends to have smaller *r_m_* and larger θ than in obligate sexuals. In cyclic parthenogens, **G** has similar but more variable *r_m_* and larger θ than in obligate sexuals.

### The paradox of cyclic parthenogenesis

The present work adds to the growing body of theoretical and observational studies that have supported the evolutionary equivalency or advantage of cyclic parthenogens over obligate sexuals (e.g. Green and Noakes 1995; Yamauchi 1999, but see Peck and Waxman 2000; Gerber and Kokko 2016). Yet paradoxically, in animals, cyclic parthenogenesis is rare compared to obligate sexual reproduction; in fact, it is known to occur in only a few clades of rotifers, crustaceans, trematodes, and insects (Herbert 1987). Why is it rare? One hypothesis is that there are steep molecular constraints against it (Herbert 1987), but to our knowledge no specific constraints have been identified. Ecological hypothesis might also be made, although they have yet to be articulated. In several species, cyclic parthenogenetic genotypes are often found in allopatry or sympatry with obligately sexual or parthenogenetic genotypes (Lynch and Gabriel 1983; Jaron et al. 2022; Gavrilov-Zimin 2024). This suggests that cyclic parthenogenesis might not be evolutionarily stable. Indeed, some theoretical work demonstrates that the benefit of occasional sex and recombination can diminish when the odds of sexual reproduction is low or if there is sexual conflict (Peck and Waxman 2000; Gerber and Kokko 2016), and observational studies indicate that cyclic parthenogenesis may often be a transition state between obligate sexuality and strict parthenogenesis (Larose et al. 2023). If any amount of parthenogenesis becomes an option, its short-term benefits are so strong that moderation might be hard to sustain. Our model points to another possibility: in cyclical parthenogens selection on **G** and **M** is less efficient than in obligate sexuals. Under the specific phenotype structures and adaptive landscapes we examined, this inefficiency does cause cyclical parthenogens to be less evolvable than obligate sexuals. But with other phenotypes and selective landscapes it could be.

## Conclusion

With individual-based models, we have shown that cyclic parthenogenesis facilitates the crossing of valleys in univariate and bivariate fitness landscapes. Moreover, we have found support for the specific mechanistic hypothesis that the crossing of fitness valleys is primarily facilitated by clonal amplification, that is, by reducing recombination during the establishment of escape genotypes. By contrast, our analysis provides weaker support for the evolutionary capacitance hypothesis that the accumulation of cryptic genetic diversity in cyclic parthenogens increases the rate at which escape genotypes are initially produced. In bivariate fitness landscapes, the size of the advantage of cyclic parthenogens in valley crossing depends on the shape of fitness valley; the evolvability gap shrinks when the leading axis of genetic diversity aligns with the global fitness optimum. With functional epitasis, cyclic parthenogenesis reduces in the alignment between their mutational variance-covariance structure and the selection surface. More work is required to resolve the paradox of cyclic parthenogens’ evolutionary advantages and rarity in animals.

## Author contribution

Z.Y. and N.B.H. designed the research, analyzed data and wrote the paper. Z.Y. performed the research.

## Acknowledgment

This paper has benefited from discussions with Matthew Wolak. We acknowledge the use of ChatGPT, a language model developed by OpenAI, for minor suggestions with respect to the texts and code.

## Data availability

Codes and data underlying this article are available in a Zenodo repository: https://zenodo.org/records/20642543.

## Reference

Arnold, S. J. 2023. Evolution of the G-matrix on a stationary adaptive landscape. Pp. 289–309 *in* Evolutionary Quantitative Genetics. Oxford University Press, Incorporated, Oxford, UNITED KINGDOM.

Bergman, A., and M. L. Siegal. 2003. Evolutionary capacitance as a general feature of complex gene networks. Nature 424:549–552. Nature Publishing Group.

Bulmer, M. G. 1974. Linkage disequilibrium and genetic variability. Genetical Research 23:281– 289.

D’Souza, T. G., and N. K. Michiels. 2010. The Costs and Benefits of occasional sex: theoretical predictions and a case study. Journal of Heredity 101:S34–S41.

Falconer, D., and T. Macky. 1996. Heritability. Pp. 160–183 in Introduction to quantitative genetics. Pearson Education Limited, Essex, England.

Gavrilov-Zimin, I. A. 2024. Gallophilous theory of cyclical parthenogenesis in aphids (Homoptera, Aphidinea). Comparative Cytogenetics 18:247–276.

Gerber, N., and H. Kokko. 2016. Sexual conflict and the evolution of asexuality at low population densities. Proceedings of the Royal Society B: Biological Sciences 283:20161280.

Green, R. F., and D. L. G. Noakes. 1995. Is a little bit of sex as good as a lot? Journal of Theoretical Biology 174:87–96.

Guillaume, F., and M. C. Whitlock. 2007. Effects of migration on the genetic covariance matrix. Evolution 61:2398–2409.

Haller, B. C., P. L. Ralph, and P. W. Messer. 2026. SLiM 5: Eco-evolutionary simulations across multiple chromosomes and full genomes. Molecular Biology and Evolution 43:msaf313.

Herbert, P. D. H. 1987. Genotypic characteristics of cyclic parthenogens and their obligate asexual derivatives. Pp. 175–194 in The evolution of sex and its consequences. Birkhäuser Verlag, Boston.

Jaron, K. S., D. J. Parker, Y. Anselmetti, P. Tran Van, J. Bast, Z. Dumas, E. Figuet, C. M. François, K. Hayward, V. Rossier, P. Simion, M. Robinson-Rechavi, N. Galtier, and T. Schwander. 2022. Convergent consequences of parthenogenesis on stick insect genomes. Science Advances 8:eabg3842. American Association for the Advancement of Science.

Jones, A. G., S. J. Arnold, and R. Bürger. 2007. The mutation matrix and the evolution of evolvability. Evolution 61:727–745.

Jones, A. G., R. Bürger, and S. J. Arnold. 2014. Epistasis and natural selection shape the mutational architecture of complex traits. Nature Communications 5:3709. Nature Publishing Group.

Kim, Y. 2007. Rate of adaptive peak shifts with partial genetic robustness. Evolution 61:1847–1856.

Lande, R. 1975. The maintenance of genetic variability by mutation in a polygenic character with linked loci. Genetics Research 26:221–235.

Larose, C., G. Lavanchy, S. Freitas, D. J. Parker, and T. Schwander. 2023. Facultative parthenogenesis: a transient state in transitions between sex and obligate asexuality in stick insects? Peer Community Journal 3.

Lynch, M., and W. Gabriel. 1983. Phenotypic evolution and parthenogenesis. The American Naturalist 122:745–764. The University of Chicago Press.

Moore, F. B. G., and S. J. Tonsor. 1994. A simulation of Wright’s shifting-balance process: migration and the thress phases. Evolution 48:69–80.

Otto, S. P. 2009. The Evolutionary Enigma of Sex. The American Naturalist 174:S1–S14. The University of Chicago Press.

Paaby, A. B., and G. Gibson. 2016. Cryptic genetic variation in evolutionary developmental genetics. Biology 5:28. Multidisciplinary Digital Publishing Institute.

Peck and Waxman. 2000. What’s wrong with a little sex? Journal of Evolutionary Biology 13:63–69.

Phillips, P. C., M. C. Whitlock, and K. Fowler. 2001. Inbreeding changes the shape of the genetic covariance matrix in Drosophila melanogaster. Genetics 158:1137–1145.

R Core Team. 2025. R: a language and environment for statistical computing. R Foundation for Statistical Computing, Vienna, Austria.

Schluter, D. 1996. Adaptive radiation along genetic lines of least resistance. Evolution 50:1766– 1774.

Steppan, S. J., P. C. Phillips, and D. Houle. 2002. Comparative quantitative genetics: evolution of the G matrix. Trends in Ecology & Evolution 17:320–327. Elsevier.

Trotter, M. V., D. B. Weissman, G. I. Peterson, K. M. Peck, and J. Masel. 2014. Cryptic genetic variation can make “irreducible complexity” a common mode of adaptation in sexual populations. Evolution 68:3357–3367.

Wade, M. J., R. G. Winther, A. F. Agrawal, and C. J. Goodnight. 2001. Alternative definitions of epistasis: dependence and interaction. Trends in Ecology & Evolution 16:498–504. Elsevier.

Weinreich, D. M., and L. Chao. 2005. Rapid evolutionary escape by large populations from local fitness peaks is likely in nature. Evolution 59:1175–1182.

Weinreich, D. M., R. A. Watson, and L. Chao. 2005. Perspective: sign epistasis and genetic constraint on evolutionary trejectories. Evolution 59:1165–1174.

Weissman, D. B., M. W. Feldman, and D. S. Fisher. 2010. The rate of fitness-valley crossing in sexual populations. Genetics 186:1389–1410.

Wright, S. 1932. The roles of mutation, inbreeding, crossbreeding and selection in evolution. Pp. 356–366 *in* Proceedings of the Sixth International Congress of Genetics. Genetics Society of America.

Yamauchi, A. 1999. Evolution of cyclic sexual reproduction under host–parasite interactions. Journal of Theoretical Biology 201:281–291.

Zheng, J., J. L. Payne, and A. Wagner. 2019. Cryptic genetic variation accelerates evolution by opening access to diverse adaptive peaks. Science 365:347–353. American Association for the Advancement of Science.

