## Supplementary material for "Cyclic parthenogenesis helps populations cross fitness valleys": Figure S

**Abbreviations:**

| **Parameter** | **Symbol** |
| --- | --- |
| Frequency of sexual generation^†^ | *ρ* |
| Recombination rate | *R* |
| SD of additive mutaional effect | *V_m_* |
| SD of epistasis coefficient | *V_e_* |
| SD of environmental noise | *V_n_* |
| Weakness of direct selection in the sub optimum | *ω* |
| Selection correlation in the sub optimum | *r_ω_* |
| Angle between optima | *α* |

^†^When *ρ* = 1, sexual reproduction is obligate. Unless otherwise mentioned, cyclic parthenogens has a *ρ* = 10.

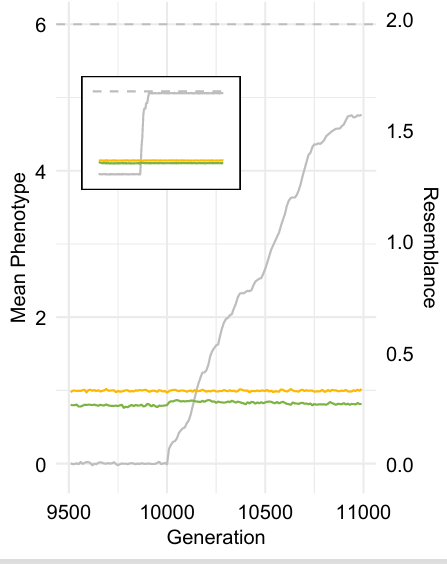

**Figure S1. Parent-offspring** genetic resemblance changes at the beginning of fitness valley-crossing in obligate sexuals. Data are measurements every 50 generations, averaged across 30 replicates under the univariate phenotype model and core parameter set. Dashed line: the global fitness optimum. Green: the resemblance between parents and offspring (*h^2^*) of the sum of genetic values; yellow: the resemblance between parents and offspring (|*h^2^*|) of the sum of absolute genetic values; grey: mean phenotype. The global fitness optimum is added at generation 10, 000. The embedded panel shows the long-term dynamic from the beginning of simulation to the 3,000^th^ generation. Only immediately after the addition of the global fitness optimum and around the onset of mean phenotype increases, *h^2^* increases while |*h^2^*| stays flight.

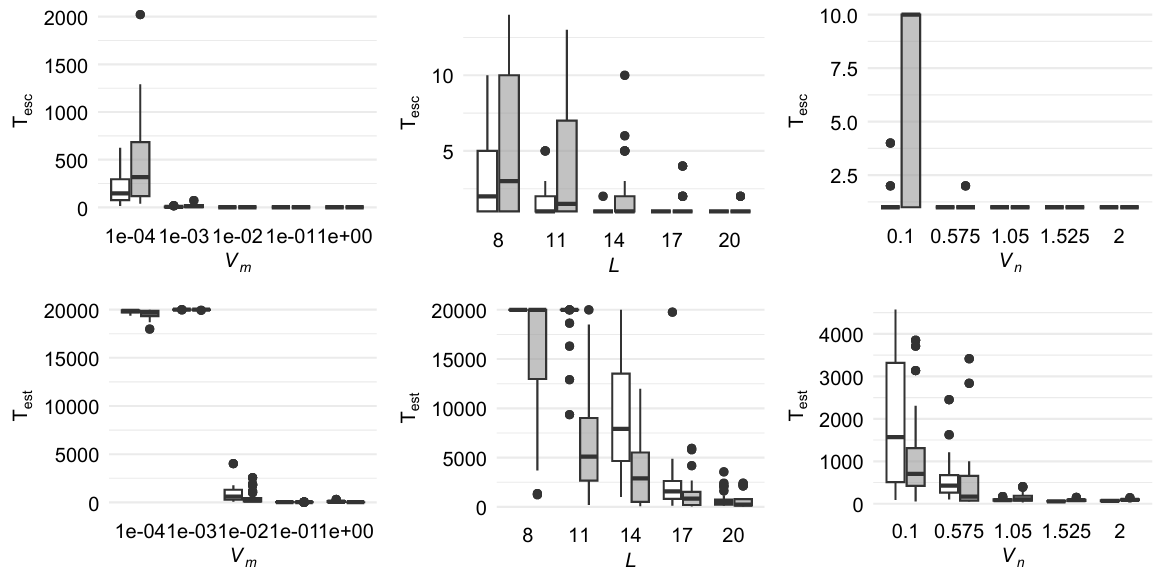

**Figure S2. Sensitivity analysis of T_esc_ and T_est_ in the univariate phenotype model.** White: obligate sexuals; grey: cyclic parthenogens. The bottom, center, and top line of each box mark the 1^st^, 2^nd^, and 3^rd^ quantiles respectively. Whiskers show 1.5 times the inter quantile distance. Each bar is from 30 replicates with the core parameter set except the parameter with the value as specified. Cyclic parthenogens have larger T_esc_ but lower T_est_ than obligate sexuals, unless the process occurs in an instant.

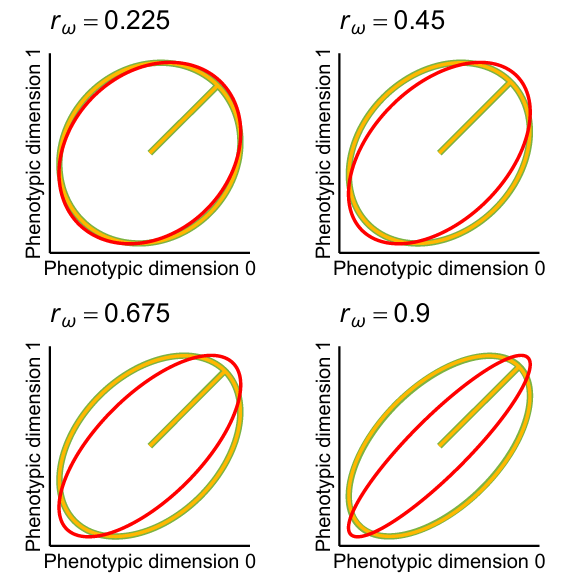

**Figure S3. 95% Confidence Ellipse of G in the bivariate, additive phenotype model.** Yellow: obligate sexual; green: cyclic parthenogens; red: the selection matrix **Ω**. Each ellipse is the average of 30 replicates. All **G**s are shaped by the corresponding **Ω** with little effect from reproductive mode.

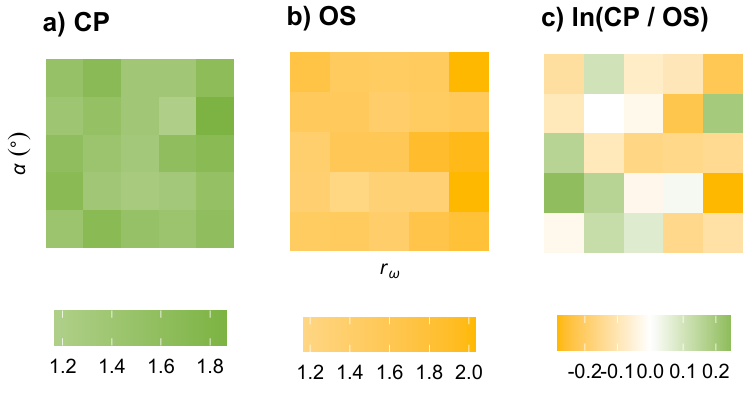

**Figure S4. T_esc_ in fitness landscapes of various shapes in a) cyclic parthenogens, and b) obligate sexuals, along with c) the log-ratio of cyclic parthenogens over obligate sexuals.** Each block is the average of 30 replicates. Five evenly distanced values on [0.0, 0.9] were used for *r_ω_* and on [0, 90°] for *α*. In contrast to T_est_ (Fig. 5), *r_ω_* and *α* do not affect T_esc_ in a systematic way. This is at least partially because T_esc_ tends to be very small. Parameter sets that allow larger T_esc_ make fitness valley-crossing unlikely.

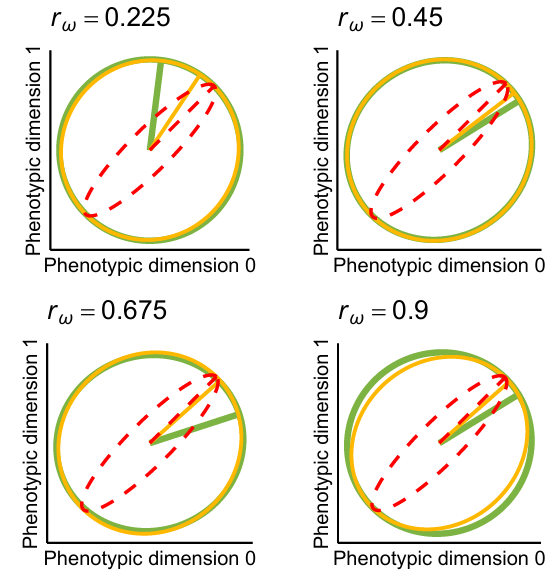

**Figure S5. 95% confident ellipses of M under the bivariate phenotype model with functional epistasis.** Each ellipse is the average of 30 replicates. Yellow: obligate sexual; green: cyclic parthenogens; red: the selection matrix **Ω**. **M** is tiny relative to **Ω**, so leading eigenvalue of all matrices are rescaled to one. **M** is shaped by **Ω** but to a more limited extent than **G** is shaped by **Ω**, as found in (Jones et al. 2014). The **M**-**Ω** alignment is stronger in obligate sexuals especially with larger *r_ω_*.

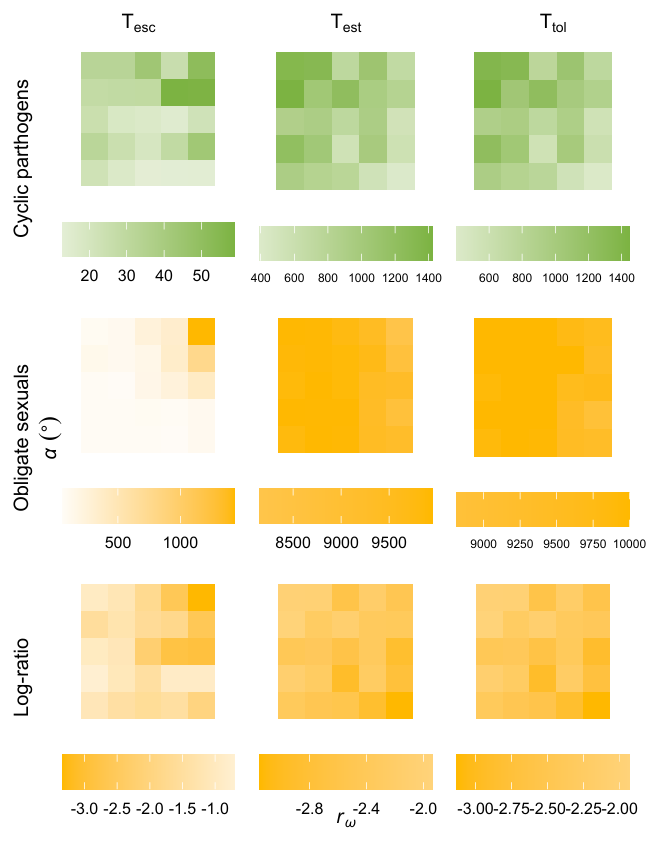

**Figure S6. T_esc_ , T_esc_ , and T_tol_ under the bivariate phenotype model with functional epistasis, with fitness landscapes of various shapes, for (top) cyclic parthenogens, and (middle) obligate sexuals, along with (bottom) the log-ratio of the former over the later.** Each block is the average of 30 replicates. Five evenly distanced values on [0.0, 0.9] were used for *r_ω_* and on [0, 90°] for *α*. For cyclic parthenogens, small *α* and small *r_ω_* minimizes T_est_ while large *α* with small *r_ω_* maximizes it. Obligate sexual T_tol_ is always larger than cyclic parthenogenetic T_est_.

**Reference**

Jones, A. G., R. Bürger, and S. J. Arnold. 2014. Epistasis and natural selection shape the mutational architecture of complex traits. Nature Communications 5:3709. Nature Publishing Group.
